# Mapping extinction risk in the global functional spectra across the tree of life

**DOI:** 10.1101/2020.06.29.179143

**Authors:** Carlos P. Carmona, Riin Tamme, Meelis Pärtel, Francesco de Bello, Sébastien Brosse, Pol Capdevila, Roy González-M., Manuela González-Suárez, Roberto Salguero-Gómez, Maribel Vásquez-Valderrama, Aurèle Toussaint

## Abstract

Although one quarter of the species of plants and vertebrates are threatened with extinction, little is known about how the potential effect of extinctions on the global diversity of ecological strategies. Using trait and phylogenetic information for more than 75,000 species of plants, mammals, birds, reptiles, amphibians and freshwater fishes, we characterized the global functional spectra of each of these groups. Mapping of extinction risk within these spectra revealed that larger species with slower pace of life are universally threatened. Simulated potential extinctions revealed extensive internal reorganizations in the global functional spectra, which are particularly severe for mammals and amphibians. Considering the disproportionate importance of the largest species for ecological processes, our results emphasize the importance of actions to prevent the extinction of the megabiota.

---

Humans have triggered a sixth mass extinction (*1*, *2*) in which nearly 1 million species are now estimated to be at risk of extinction (*3*). However, counts of threatened species do not fully reflect the ecological impacts of extinctions because species’ contributions to ecosystem functioning depend on their functional traits (*4*, *5*). Consequently, extinctions of species with unique traits are likely to have more dramatic consequences on ecosystems than extinctions of species with redundant traits (*6*). Yet, little is known about the impacts of the ongoing mass extinction on the functional diversity for the majority of organisms. Improving our understanding of the factors modulating species’ extinction risk is critical for conservation (*7*–*10*). These factors include functional traits —morphological, physiological, phenological or behavioural features that govern the functional role of organisms and the effects that environment have on them (*11*, *12*).

Variation among species in terms of traits is remarkable, encompassing differences of several orders of magnitude, such as the differences in body size between the less than 2 g of shrews and more than 100 Tn of whales in mammals, or the differences in seed mass between the less than 1 microgram of some orchids and more than 15 Kg of coconuts in plants. Despite all this variation, species’ ecological strategies resulting from trait combinations are constrained by physiological limits set by evolutionary history and trade-offs in resource allocation (*13*–*15*). Accordingly, recent mappings of the global trait spectra of plants (*16*), and birds and mammals (*17*) have described them as two-dimensional surfaces, where occupation of the trait space is restricted compared to null expectations. There are still no characterizations of the global trait spectra of other groups of vertebrates, which precludes understanding how extinction risk is distributed within these functional spaces. So far, the few characterizations of the impact of species extinctions have only considered reductions in the amount of functional space occupied by each taxonomic group (*17*–*19*). However, functional redundancy among species (*i.e.* adjacent species in the functional space; *16*, *20*) is very widespread. This means that it is likely that many of the functional consequences of extinctions do not only affect the overall volume and boundaries of the functional spectra, but deeply reorganize its internal structure. Such changes can be better examined with probabilistic approaches that consider shifts in the density of occupation of the functional space (*20*–*22*), hence fully accounting for the potential effect of redundancy between species.

Here, using traits for more than 75,000 species of plants, mammals, birds, reptiles, amphibians, and freshwater fish, we describe the impacts of the potential extinction of threatened species on functional diversity across the multicellular Tree of Life. By mapping extinction risk in the functional space of each group, we found that extinction risk is not randomly distributed, but localized in certain areas of the functional space occupied by species with large size, slow pace of life, or low fecundity. Although global losses of functional space are lower than the rates of species extinctions, we show that extinctions will lead to a denser functional aggregation of species at the global scale. Potential extinctions will therefore cause dramatic erosion and rearrangement of ecological strategies across the tree of life.

## Data and methods

We used functional trait information from over 39,260 species of plants, 4,953 mammals, 9,802 birds, 6,567 reptiles, 6,776 amphibians and 10,705 freshwater fishes from different published databases (*19*, *23*–*25*). For each of these groups, we chose a set of fundamental functional traits associated with different key aspects of their ecology (see Methods and table S1 for further details). We first characterised the global functional space occupied by each group by means of principal component analyses (PCA) based on the compiled traits (*16*, *17*).

To account for functional redundancy between organisms, we used a trait probability density approach (TPD), which allows mapping the functional spectra of organisms as probabilistic surfaces (*20*). Instead of simply characterizing the boundaries of the spectra, the TPD approach reflects the abundance of species with similar suites of traits (*16*, *21*). This method effectively represents the functional spectra as a landscape with “peaks” that reflect areas with high density of species and “valleys” where the density is lower, and allows to test changes in aspects of trait space occupation other than volume. Specifically, we examined some properties of these spectra, such as the degree of aggregation of species in particular areas of the functional space (functional hotspots; *16*) and the positioning of such areas within the occupied space.

We then used IUCN categories to map extinction risk in the functional trait space of each group by means of generalized additive models (GAM; *26*), which allowed us to discover substantial differences in extinction risk between functional strategies. Finally, we used simulated extinctions based on the IUCN species’ extinction risk assessments to define the functional spectra of a world where threatened species have gone extinct, and compared these with an alternative scenario of random species extinctions. Matching the current functional spectra with the spectra after possible extinctions allowed us to reveal the degree of “erosion” that could be potentially experienced by global functional diversity, exposing functional areas that will become particularly vulnerable to further extinctions.

## High redundancy in the global functional spectra

For five groups, the resulting functional spaces consisted of two main dimensions (*27*, *28*), which captured 68-83% of the total functional trait variation, with the only exception of freshwater fishes, whose main functional space extended over four dimensions (table S1). In all cases, the first principal component was linked to traits related to the size of the organisms, such as plant height, body length, or body mass (Fig. 1). The second principal component was frequently related to traits linked to reproduction, such as the frequency and amount of offspring produced.

**Fig 1.**
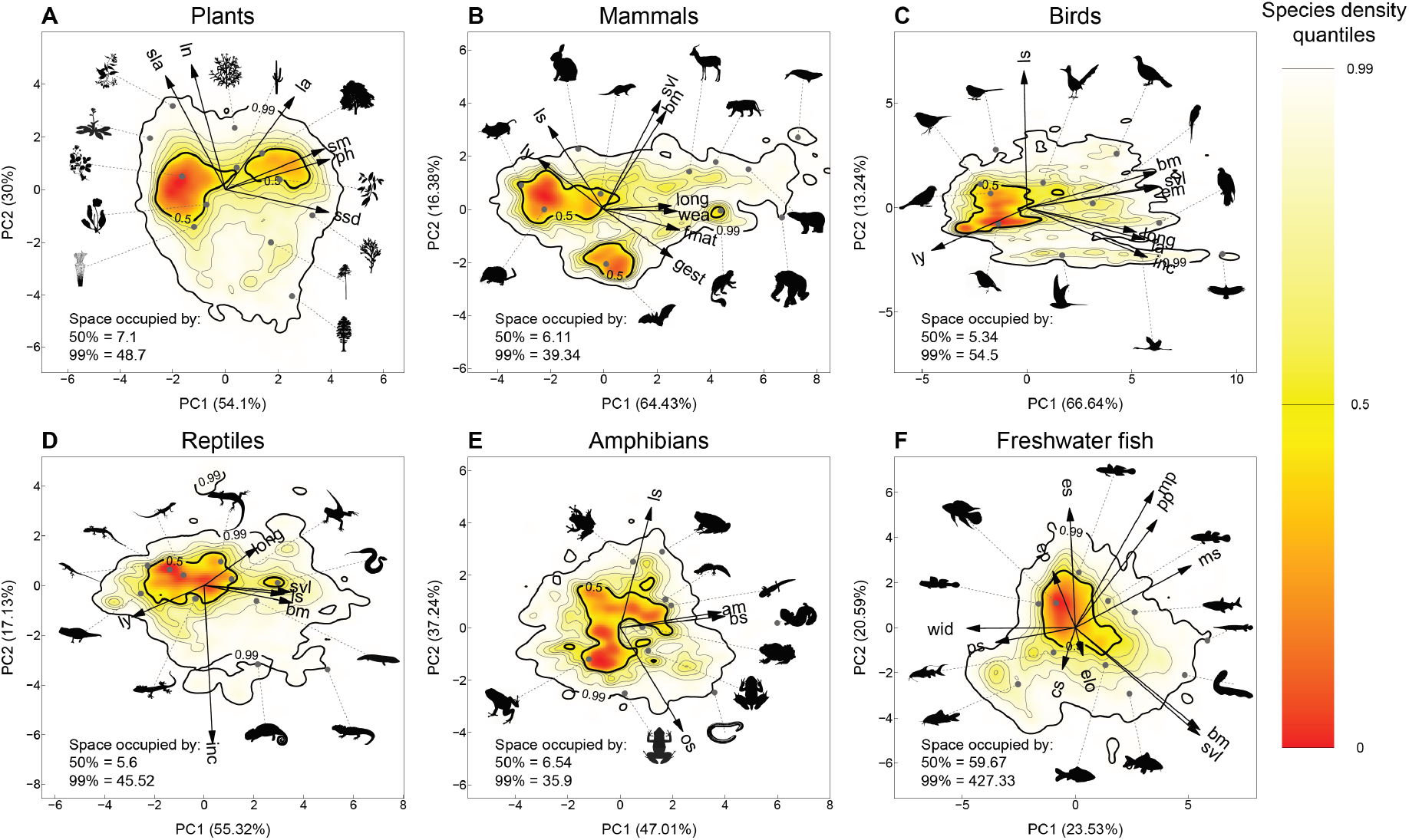
Global functional spectra of plants (A), mammals (B), birds (C), reptiles (D), amphibians (E) and freshwater fish (F). Probabilistic species distributions in the spaces defined by the two first principal components of PCA analyses (details in table S1) considering different functional traits for each group (see table S3 for definitions of each functional trait). Arrows indicate the direction and weighting of each trait in the PCA (see Methods for description of each individual trait, and table S3 for the meaning of the traits). The colour gradient (red-yellow-white) depicts different density of species in the defined space (red areas are more densely populated). Arrows show the loadings of the considered traits in the resulting PCA. Thick contour lines indicate the 0.5 (hotspots, see main text) and 0.99 quantiles, and thinner ones indicate quantiles 0.6, 0.7, 0.8 and 0.9. Silhouettes were downloaded from PhyloPic (www.phylopic.org). The legends within each panel show the amount of functional space (measured in standard deviation units, SD^2^, except for freshwater fish, in SD^4^) occupied by the hotspots (0.5 quantile) and the 0.99 quantile distribution.

Our analyses reveal high degrees of lumpiness (and hence of functional redundancy; *29*) in the occupation of all the functional spaces. As a result, the amount of functional space occupied was always much smaller than in equivalent multivariate normal distributions (fig. S1). In particular, the hotspots of the different groups (*i.e.* the smallest portion of functional space including 50% of the species; *16*) were consistently noticeable. For example, in the case of birds, half of the species occupied only 9.8% of the total spectrum, whereas amphibians were less aggregated (18.2% of the total spectrum).

The plants and mammals’ spectra displayed multiple distinct hotspots. For plants, one of the hotspots is occupied by grasses and herbs, and the other by woody species, in agreement with (*16*) (Fig. 1A). Mammals showed two main hotspots separated along the second PCA axis, one corresponding to species that reproduce often and produce many offspring with relatively short gestation times, whereas the other included species with smaller reproductive outputs and longer gestation times. The third mammal hotspot, much smaller, included primarily primate species (suborder Simiiformes) with long lifespans and late weaning times (Fig. 1B).

Birds, reptiles, amphibians and fish displayed a single hotspot, with skewed distributions of species in their size-related axes for all taxa but fishes (even after log-transformation before PCA). These spectra revealed large proportions of small species and much fewer species of larger size (Fig. 2), in accordance with previous observations (*17*, *30*). Therefore, the hotspots tended to be close to the boundaries of the occupied space, as revealed by the high values of functional divergence (fig S2). In contrast, the freshwater fish hotspot was centred within the functional space, revealing that median-sized species with generalist morphology represent the core of the freshwater fish fauna.

**Fig 2.**
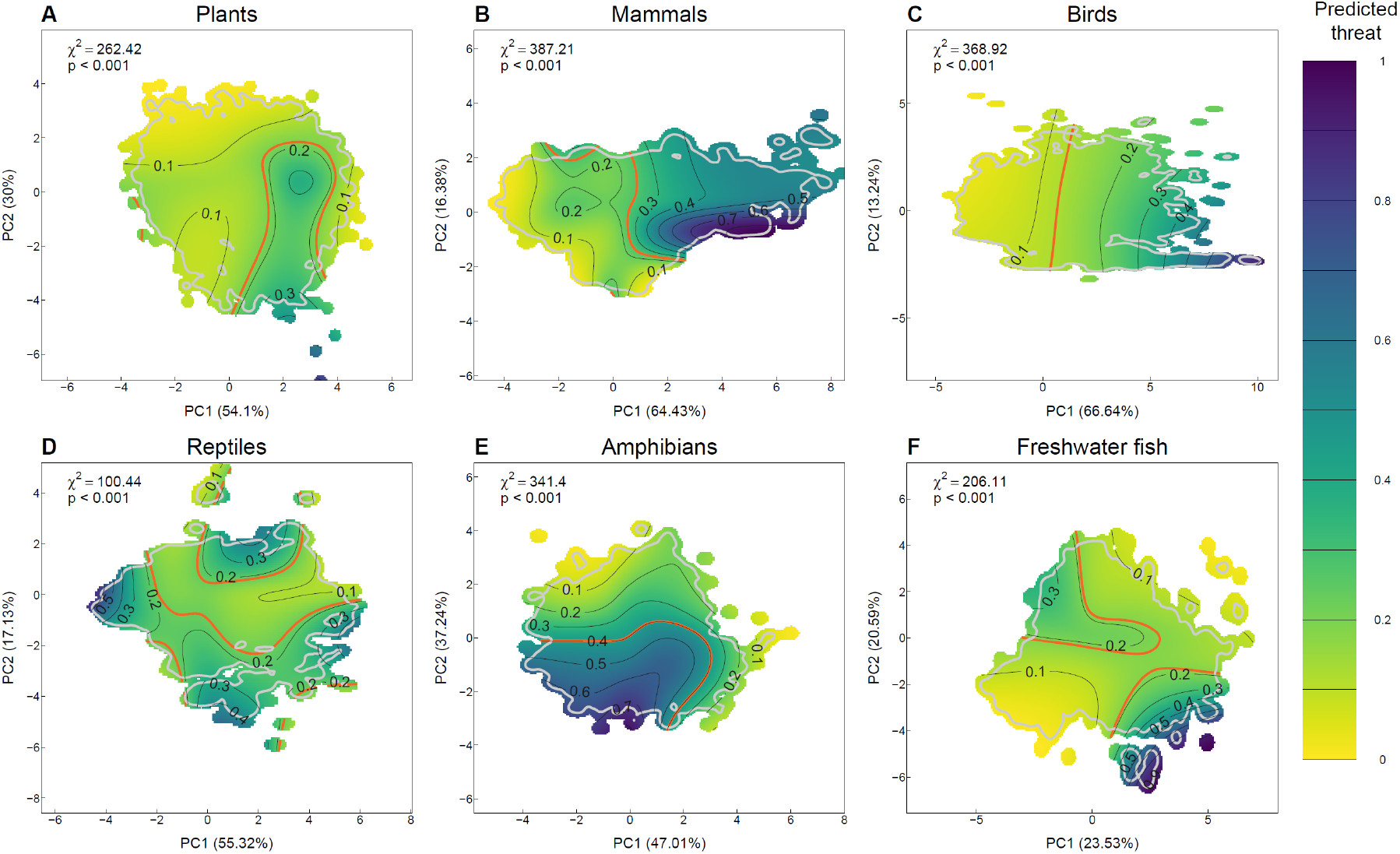
Extinction risk in the functional spaces of plants (A), mammals (B), birds (C), reptiles (D), amphibians (E) and freshwater fish (F). Probability of species being classified as threatened (see Methods) according to generalized additive models (GAM with binomial distribution) using the position of species in the functional space as predictors. Yellow tones indicate lower risk of extinction, whereas purple tones indicate high risk of extinction. Legend shows χ^2^ and p values of the GAM corresponding to each taxonomic group. For each group, the red contour lines indicate the average threat probability (proportion of species classified as threatened in the group). The grey line indicates the 0.99 quantile of the spectra of each group considering only species whose threat status is known.

## Mapping extinction risk

The position of a given species along the multivariate trait space predicts its conservation status, regardless of taxonomic group examined. Threatened species showed distinct occupation patterns of the different functional spaces, evidenced by the lower similarity between the global spectra and the spectra of threatened species than between the global spectra and the spectra of non-threatened species in all taxa (table S3). Accordingly, mapping of threat probabilities revealed substantial increases in the risk of being threatened in areas of the functional space occupied by species with larger sizes and slower or less copious reproductive outputs being much more likely to be threatened (Fig. 2 and fig. S3). In particular, for plants, the probability of being threatened was up to three times higher for woody species than for herbaceous ones (Fig. 2A). Mammal species with long weaning and gestation periods, with relatively large sizes (right end of the functional spectra) showed up to eight times higher threat risk than smaller species towards the left side of the spectrum (Figs 1B and 2B). The pattern for birds was similar, with species with later fledging ages, longer incubation times and larger sizes having up to six times higher threat risk than smaller species with faster breeding times (Figs 1C and 2C). For two groups, freshwater fishes and reptiles, in addition to large species, threat risk also increased toward species with smaller size (Fig. 2D,F).

The high degree of aggregation observed in the examined taxonomic groups acts as a buffer against extinctions. Accordingly, projected losses of functional space after extinctions were substantially lower than the proportion of threatened species, ranging between 1.53% (plants) and 13.62% (freshwater fishes; Fig. 3). Despite this finding, the proportion of lost space after extinctions was higher than expected under a random species extinction hypothesis for mammals, reptiles and freshwater fishes, revealing that the diversities of functional strategies of these groups are particularly vulnerable to extinction. Nonetheless, projected extinctions will erode the functional spectra in ways that go beyond the amount of functional space being completely lost. To show this pattern, we compared the functional spectra with and without threatened species, discovering that the spectra of all groups will experience much larger shifts than expected under random extinctions (Fig. 3). These shifts were particularly striking for some groups (movies S1-6), including mammals, in which most of the functional diversity erosion would take place close to the boundaries of the functional spectra. Thus, further extinctions would increase the risk of completely losing parts of the spectra corresponding to mammal species with high longevity, late sexual maturity, and long gestation and weaning periods. For instance, the hotspot formed by primate species is projected to disappear due to the high prevalence of threat in the species composing it (movie S2). The shift in the functional spectra of amphibians after extinctions, characterized by a marked decrease in the relative proportion of species with small reproductive outputs, was also remarkable (movie S5). For freshwater fish, although functional changes were mainly clustered toward particular boundaries of the functional space, corresponding for instance to the large sized species, that are, as for other taxonomic groups, long lived species. In general, the higher risk of extinction of large, long-lived and slow-reproducing species will result in decreased functional redundancy for these species at the global scale. Lower redundancy may in turn lead to higher vulnerability to future extinctions for the affected functional strategies.

**Fig 3.**
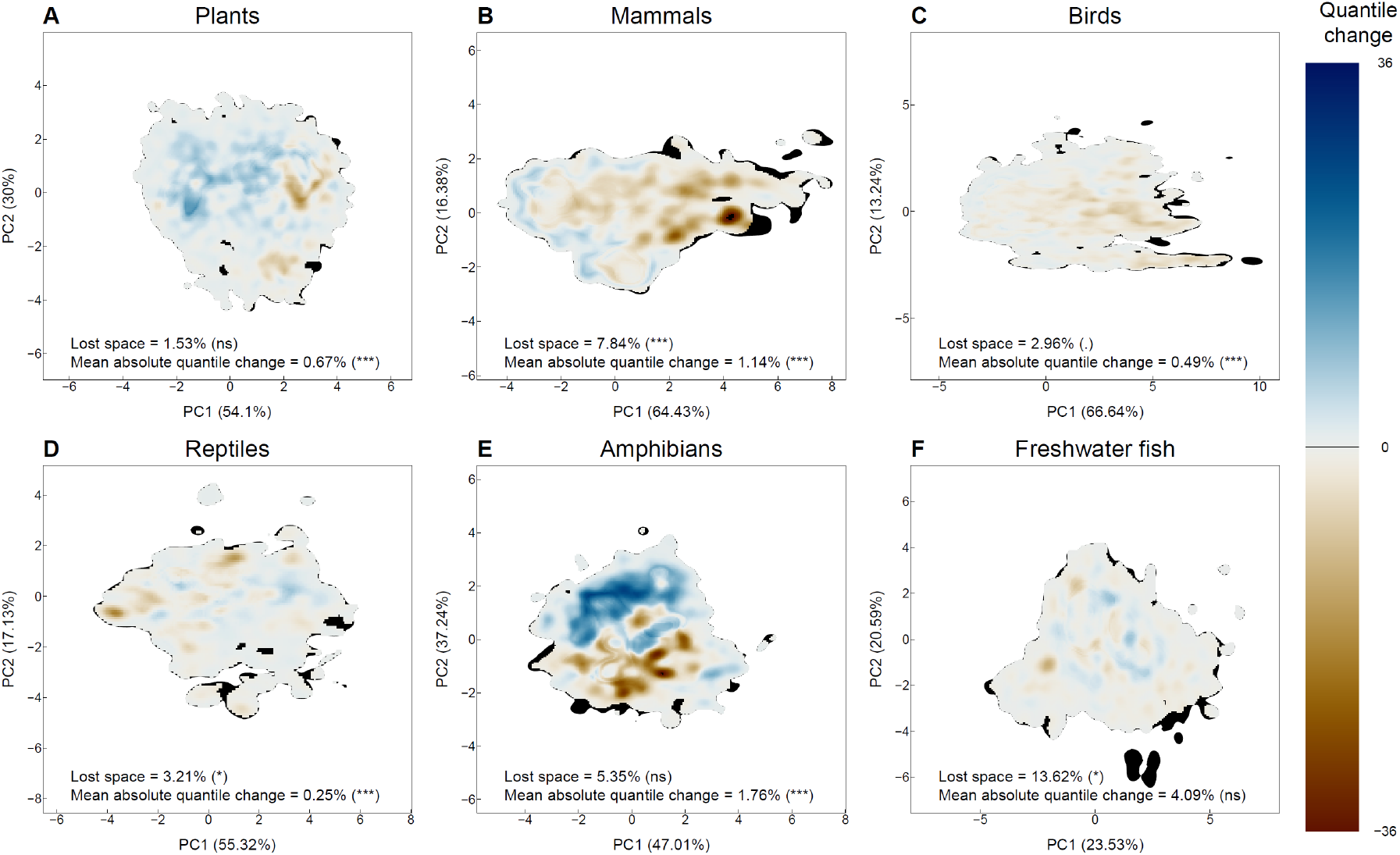
Shifts in functional spectra after simulated extinctions for plants (A), mammals (B), birds (C), reptiles (D), amphibians (E) and freshwater fish (F). Differences (expressed in quantile changes) between the functional spectra of species assessed by IUCN before and after removing species classified as threatened. Brown tones reflect areas in which the TPD quantiles are lower after extinctions (i.e. those traits become relatively less frequent at the global scale), and blue tones reflect areas in which the TPD quantiles increase (i.e. those traits become relatively more frequent at the global scale). Black areas show the parts of the functional space that would disappear from the global spectra after extinctions (trait combinations that go completely extinct). The legend for each group shows the proportion of total space that would be lost after extinction (expressed as percentage) and the average absolute change in quantiles across the spectra; p-values come from comparison with a null model (1,000 repetitions) in which the same number of species are lost at random. P-values: * P<0.05, *** P<0.001, ns P>0.05. Significant values always indicate higher values (either lost space or quantile change) than expected by chance.

## Discussion

Using the most species diverse trait databases collected to date, we show that, in most groups, realised functional strategies are constrained to a single plane in which species are clumped around a few strategies that are rather prevalent (*16*, *17*, *31*). Contemporary functional spectra are probably rather different from those in the relatively recent past due to extinctions in the last millennia (*32*). For example, since the Pleistocene, larger animal species have gone extinct at much higher rates than smaller species (*33*). This trend continues nowadays, triggered by the rise of human population and technologies that disfavours large and slow pace of life species (*9*, *33*, *34*). In addition, the smallest animal species are also among the most threatened (*35*), as already reported for fish and reptiles, because small animals often have low dispersal abilities and remain endemic from restricted areas, making them vulnerable to extinction (*7*, *10*). Our models show that large-sized, slow-paced and slow reproducing species are much more likely to go extinct across all groups, and provide some support for the notion that small species are also more threatened in the case of reptiles and freshwater fishes. Mammals and amphibians emerge as the groups most affected by these changes, experiencing shifts in the functional space towards faster-living and highly-reproductive strategies, respectively. These results are in line with the notion that larger organisms are more sensitive to global change, a trend affecting plants, land and aquatic animals that will probably be exacerbated in the future (*32*, *36*, *37*).

We show that estimated extinctions will cause noticeable reorganizations in the functional spectra of most groups, making them more vulnerable to further extinctions in the future. Although the impacts of extinctions on multivariate functional diversity is increasingly being studied, research focus has so far been on quantifying losses in the amount of functional space occupied (*17*–*19*). However, functional diversity encompasses all aspects of trait variation among organisms, of which total occupation (functional richness; *38*) is only one component (*21*, *38*). Here, we show that species extinction impacts on functional richness alone appear to be much milder than the intense erosion observed when the whole spectra is considered. It is important to note that extinction effects in the global spectra are likely even greater in local communities. For example, although extinction risk is much higher for tree species, the global spectrum of plants is not likely to experience dramatic shifts due to high functional redundancy. However, in local communities with fewer species, functioning could be dramatically altered even by the extinction of a low proportion of species, and amplified by cascades of secondary extinctions (*39*, *40*). Such impacts are likely to be particularly important if the lost species are the largest ones, given their disproportionate importance for ecological processes (*41*). Understanding how the effects of extinctions downscale from the global scale into smaller scales (continents, realms, regions, local communities) emerges as an important priority to understand the impacts of extinction on ecosystem functioning.

## Acknowledgments

The study has been supported by the TRY initiative on plant traits (http://www.try-db.org). The TRY initiative and database is hosted, developed and maintained by J. Kattge and G.Boenisch (Max Planck Institute for Biogeochemistry, Jena, Germany). TRY is currently supported by Future Earth/bioDISCOVERY and the German Centre for Integrative Biodiversity Research (iDiv) Halle-Jena-Leipzig.

## Author contributions

CPC, AT, RT, MV-V and RG-M conceived the study. CPC, AT and RT integrated and cleaned data. SB and AT contributed data. CPC and AT designed and performed analyses. CPC wrote a first draft of the manuscript, and all authors contributed to interpret and analyse results, and contributed to revisions.

## Supplementary Materials

### Materials and Methods

#### Data collection and processing

##### Functional traits and phylogenies

We collected published information on functional traits for all the studied groups of organisms from different sources (see table S1 for detailed descriptions of each trait):

###### • Vascular plants

We used six traits previously shown to capture global spectrum of plant form and function (*16*): plant height (ph, m), specific stem density (ssd, g/m^3^), leaf area (la, mm^2^), specific leaf area (sla, mm^2^/mg), N content per unit leaf mass (ln, mg/g) and seed mass (sm, mg). We used publicly available data for these traits from the latest version of the TRY Plant Trait Database (version 5.0, https://www.try-db.org/TryWeb/Home.php, accessed April 2019; *23*). Altogether our dataset included over 955,000 trait measurements for 44,431 vascular plant taxa. In the analysis, each taxon was represented by an average trait value (excluding outliers with >3 SD). To account for within-species variation, the averages for each species-trait combination were calculated first within individuals (if multiple measurements were taken from a single individual), then within datasets (if multiple individuals were measured in the same location) and finally within species (if multiple individuals were measured in various locations).

Plant height data included 179,263 measurements of adult plant vegetative height for 15,008 taxa. In most datasets this was represented as observed height or average of measurements. In some cases, plant height was represented as the maximum observation (12,942 records). Specific stem density (SSD) data included 28,723 measurements for 8,840 taxa. As this trait is usually measured for woody species, we estimated SSD for herbaceous plants using leaf dry mass content information (127,067 measurements for 5,764 taxa), following the procedures described in (*16*). Leaf area data included 119,172 measurements for 13,928 taxa. Different datasets in TRY reported various measurements of leaf area (e.g. leaflet or leaf, petiole included or excluded). To maximize our data coverage, we included both leaflet and leaf measurements, and preferred measurements including petiole (if both data types, petiole included or excluded, were reported for the same individual). Specific leaf area data included 223,126 measurements for 10,674 taxa. Similarly to leaf area data, we preferred measurements that included petiole. Data for nitrogen (N) content per unit leaf mass included 92,850 measurements for 10,530 taxa. Data for seed mass included 185,182 measurements for 25,394 taxa.

###### • Mammals, Birds and Reptiles

We used the Amniote database (*25*) including data for 4,953 species of mammals, 9,802 species of birds and 6,567 species of reptiles. The database includes data for 29 traits, but information is very incomplete for many of them (see table S3). Hence, for each group, we selected subsets of traits with sufficient information. In the case of mammals, we selected eight traits: litter size (ls, number of offspring), number of litters per year (ly), adult body mass (bm, g), longevity (long, years), gestation length (gest, days), weaning length (wea, days), time to reach female maturity (fmat, days), and snout-vent length (svl, cm). We also selected a total of eight traits for birds: clutch size (ls, number of eggs), number of clutches per year (ly), adult body mass (bm, g), incubation time (inc, days), longevity (long, years), fledging age (fa, days), egg mass (em, g) and snout-vent length (svl, cm). Finally, we selected six traits for reptiles: clutch size (ls, number of eggs), number of clutches per year (ly), adult body mass (bm, g), incubation time (inc, days), longevity (long, years) and snout-vent length (svl, cm).

###### • Amphibians

We used the AmphiBIO database (*24*) to get data for 6,776 species of amphibians (see table S3). Within this dataset, we selected a total of four traits with sufficient information: age at maturity (am, years), body size, measured in Anura as snout-vent length, and in Gymnophiona and Caudata as total length (bs, mm), maximum litter size (ls, number of individuals), and offspring size (os, mm).

###### • Freshwater fishes

We used the last updated version of the most comprehensive database to get morphological traits, available for 10,705 species of strictly freshwater fishes (*19*, *42*). Morphological measures and conventions are detailed in previous studies (*19*, *42*, *43*). All morphological traits have been measured on side view picture, using one specimen per species and applying conventional rules for unusual morphologies (e.g. species without tail, flatfishes) as defined in previous studies (see details in *19*, *43*). Briefly, this database encompasses 11 traits describing size and shape body parts involved in food acquisition and locomotion. The fish body shape and weight was described through the size using the standard length (svl) and body mass (bm) taken directly from FishBase (*45*), body elongation (elo, ratio between body length and body depth) and body lateral shape (bls, ratio between head depth and body depth). The other traits describing the position and the size of each part of the fish were: eye size (es) and position (ep), mouth size (ms) and position (mp), pectoral fin size (ps) and position (pp), and caudal peduncle throttling (cs).

###### • Phylogenies

We obtained published phylogenies for each of the considered groups (*46*–*52*). Species that were not present in the phylogeny were added to the root of their genus, using the ‘add.species.to.genus’ from the R package Phytools (*53*). Species for which we did not have evolutionary information were removed from the trait databases before the missing trait imputation procedure (see below).

##### Conservation status of species

We collected conservation status of species from the IUCN Red List (*54*) (retrieved 25^th^ September 2019) using the R package ‘rredlist’(*55*). We reclassified the IUCN categories as ‘Threatened’ (including the “Extinct in the wild”, “Critically endangered”, “Endangered”, and “Vulnerable” categories) or ‘Non-threatened’ (including the “Near Threatened” and “Least concern” categories).

##### Taxonomic standardization

Taxonomies from all the used sources (trait databases, phylogenies and IUCN Red List), were standardized using the R packages taxize (*56*) (for animals) and Taxonstand (*57*) (for plants). In the case of animals, all names were resolved against the GBIF Backbone Taxonomy (*58*), whereas in the case of plants we used The Plant List (*59*).

##### Imputation of missing traits

None of the functional trait databases assembled were complete (see table S3). We completed this information by performing a trait-imputation procedure for each group using the missForest R package (*18*, *60*). Before the imputation process, all traits were log-10 transformed, centred and scaled. The missForest package uses random forest techniques to impute trait data, which allows to include phylogenetic information in the trait imputation process, which is known to improve the estimations of missing values (*61*). We included the evolutionary relationships between species in the imputation process by including the first ten phylogenetic eigenvectors in the matrix to be imputed, as recommended in (*61*). The final numbers of species with functional trait information used in each group are shown in table S3.

#### Construction of the global spectra

We identified the main axes of functional trait variation by performing principal component analyses (PCA) on the log-transformed and scaled functional traits of each group. We used Horn’s parallel analysis in the R package paran (*28*) to determine the number of axes retained in these PCA; we will refer to these reduced spaces as functional spaces from now on. We checked the reliability of the functional spaces obtained with imputed functional trait values by comparing them with the spaces that were based only on species with complete functional information. We performed this comparison by estimating the correlation between distance matrices of the species that were common to the two spaces (space with imputed data and space with complete species only) through a Procrustes of each taxonomic group (*16*), using the ‘procuste.rtest’ function from the R package ade4 (*62*). To assess the significance of the correlation, permutation tests (9,999 randomizations) based on Monte-Carlo simulations were generated. All the Procrustes tests were highly significant (P=0.0001 in all cases, see table S2), indicating a strong correspondence between the complete and imputed functional spaces; consequently, we used the PCA based on imputed trait data in the rest of analyses.

We estimated the probabilistic distribution of the species within the functional spaces by performing multivariate kernel density estimations with the ‘TPD’ and ‘ks’ R packages (*20*, *29*, *63*, *64*). The kernel for each species was a multivariate normal distribution centered in the coordinates of the species in the life history or functional space and bandwidth chosen using unconstrained bandwidth selectors from the ‘Hpi’ function in the ‘ks’ package (*63*–*65*). The aggregated kernels for all species in a group result into the trait probability density function (TPD; *20*, *21*, *65*) of that group in the corresponding space. Although TPD functions are continuous, in order to perform operations with them it is more practical to divide the functional space into a D-dimensional grid composed of many equal sized cells (we divided the 2-dimensional spaces in 40,000 cells, 200 per dimension, and the 4-dimensional space in 810,000 cells, 30 per dimension). Then, the value of the TPD function is estimated for each cell. The value of the ‘TPD’ function in a given point of the space reflects the density of species in that particular area of the space (i.e. species with similar functional traits). For each of these spaces, we represented graphically the global TPD as well as the contours containing 50%, 60%, 70%, 80%, 90% and 99% of the total probability.

We compared the distribution of species within the different functional spaces with a null model considering that species are distributed following a multivariate normal distribution (*16*). For this, for each taxonomic group, we drew 199 samples of 1,000 simulated species from multivariate normal distributions with the same mean and covariance matrix as the observed spectra. For each of these samples we estimated a TPD function and measured functional richness (amount of space occupied by the spectra (*20*, *21*, *38*, *67*)) at the 99% and 50% quantile thresholds, and functional divergence (which indicates the degree to which the density of species in the functional trait space is distributed toward the extremes of the spectra; *20*, *38*). Then, we drew 199 samples of 1,000 species from the observed global species pool of each taxonomic group and performed similar analyses. We compared the estimations of functional richness (at 50% and 99% quantile thresholds) and functional divergence of the observed and simulated data by means of two-tailed t-tests.

#### Effects of extinctions on global functional diversity

We estimated TPD functions in the different functional spaces for the sets of species classified as Threatened or Non-threatened, as well as the TPD functions for all the species assessed by IUCN (including both threatened and non-threatened), using the same procedure described above. We then estimated the similarity between these TPD functions and the global spectra (the global TPD function considering also species not assessed by IUCN) as the overlap between the TPD functions (*21*, *68*–*71*). Compared to methods that consider exclusively the boundaries of the distributions (e.g. hypervolumes or convex hulls (*17*, *18*, *72*–*74*)), TPD-based probabilistic overlap considers also the differences in density within those boundaries. This approach provides a more complete idea of what the differences between the functional spectra are, particularly in cases where functional redundancy is high (*71*, *75*–*77*). Given that a high proportion of the considered species might be clumped in particular areas of the considered space (*16*, *17*), this methodological aspect can be particularly useful to detect differences in the occupation of functional spaces between groups of species with different conservation status. Estimating the similarity between the different groups allowed us to examine: 1) whether there is any bias regarding which species have been assessed by IUCN (overlap between the global distribution and the IUCN TPD functions), 2) whether non-threatened and threatened species occupy the considered space in different ways (overlap between the non-threatened and the threatened TPD functions). In all vertebrate taxa, the species whose conservation status has been assessed by IUCN formed a relatively random subset of the functional spectra of the different groups (high similarity between the assessed species and the global spectra; table S3). Plants constituted an exception to this pattern, reflecting the bias towards trees in the IUCN Red List (*78*) (fig. S2), as well as the high proportion of species for which conservation status is not known (in our dataset, roughly 80% of the plant species with trait measurements have not been assessed or are classified as data deficient).

After examining the overlap of TPD functions, we mapped the conservation status of species within the functional spaces. For this, we considered only species assessed by IUCN. The relationship between conservation status (1: Threatened; 0: Non-threatened) and the position in the corresponding space (PCA axes) was analyzed using a tensor product smoother-based generalized additive model (GAM; *26*) with a binomial response (R package ‘mgcv’; *78*). We then mapped the predictions of the models (including the 95% confidence intervals of the means) to visually examine how different combinations of functional traits affect the probability of species being threatened.

Finally, we explored the potential effects of the loss of threatened species on the distribution of species in the functional spaces. For this, in each space, we estimated a TPD function considering all the species assessed by IUCN, and another TPD function after removing the species classified as threatened. Whereas the TPD functions of all species assessed by IUCN reflect the current spectra, the TPD functions after removing threatened species reflect the potential spectra if all threatened species go extinct. TPD functions are probability density functions, so that they integrate to 1 across the whole functional space. We applied a quantile threshold of 99% to reduce the potential effect of outliers on the estimation of the amount of functional space occupied by the different spectra (*20*, *66*, *73*). After thresholding, the TPD functions were rescaled, so that they again integrated to 1 across the functional space, and the probabilities expressed in terms of quantiles to ease interpretability of the results.

We represented the impact of simulated extinctions by subtracting, in each cell, the quantile value of the TPD function after removing threatened species from the quantile value of TPD function of IUCN-assessed species. Negative values in this index indicate a decrease in the relative abundance of the trait values corresponding to that cell, and vice versa. To quantify how much the functional spectra of each group will change after extinctions, we estimated for each cell the absolute value of this quantile difference, and averaged these values across cells (mean absolute quantile change). With this approach we could also characterize which cells become empty after extinctions (lost space; expressed as a proportion of the total space occupied by the IUCN-assessed species spectra). While mean absolute quantile changes reflect the sensibility of the whole spectra to extinctions, lost space quantifies the amount of the functional spectra that go globally extinct. To assess whether changes in these indices were bigger than expected under a similar number of random extinctions, we also created 1,000 TPD functions simulating cases in which the same number of species were lost at random from the total set of IUCN-assessed species (i.e. including both non-threatened and threatened species rather than only threatened ones). We performed similar comparisons (i.e. lost space and mean absolute quantile change) between each of these simulated spectra and the spectra of the IUCN-assessed species, and estimated a p-value for each index and group of species by ranking the observed changes among the simulated ones. This strategy allowed us to ascertain whether losing threatened species affects reduces more or less than expected the functional spectra of the different groups (in terms of lost space and mean absolute quantile changes).

**Table S1.**
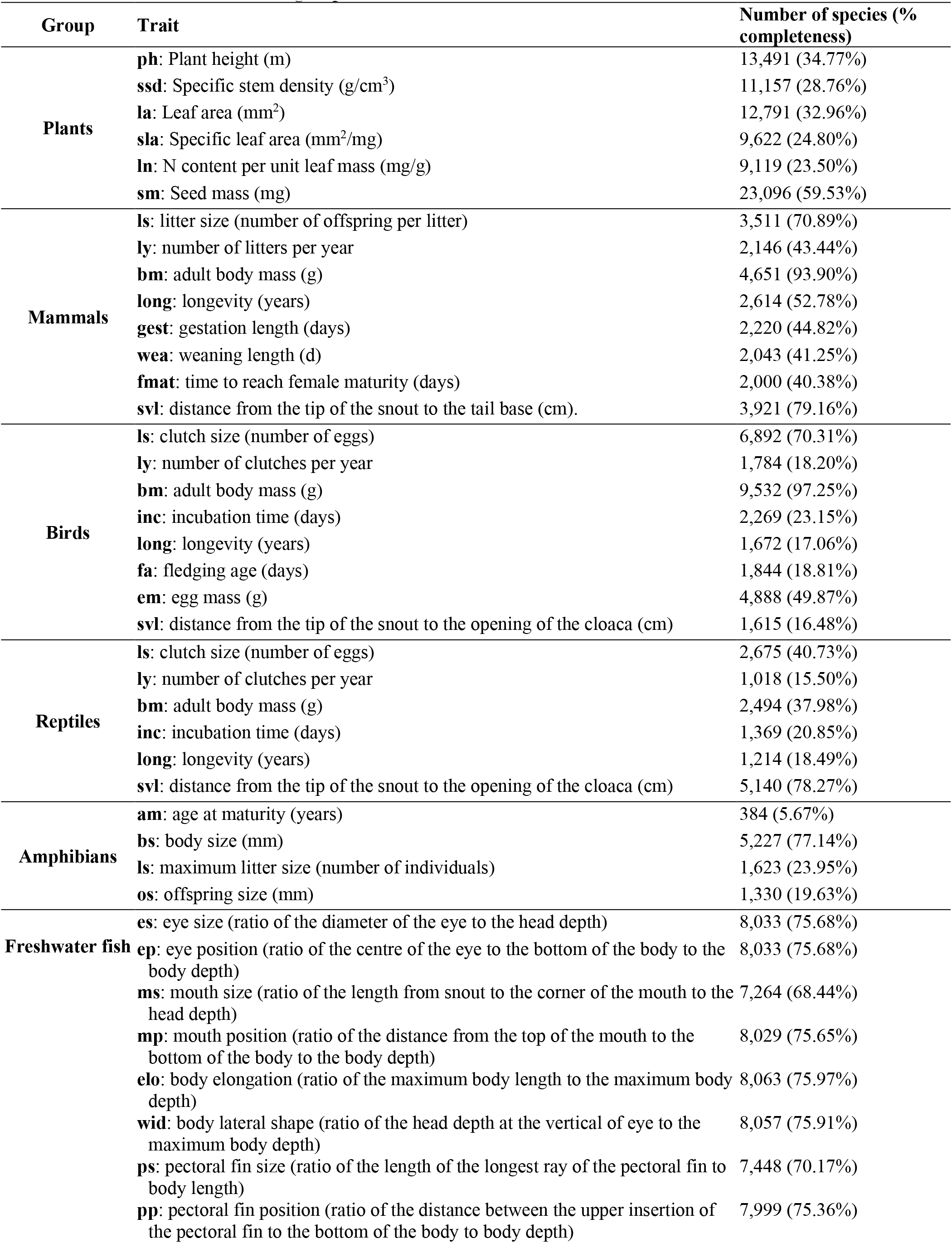
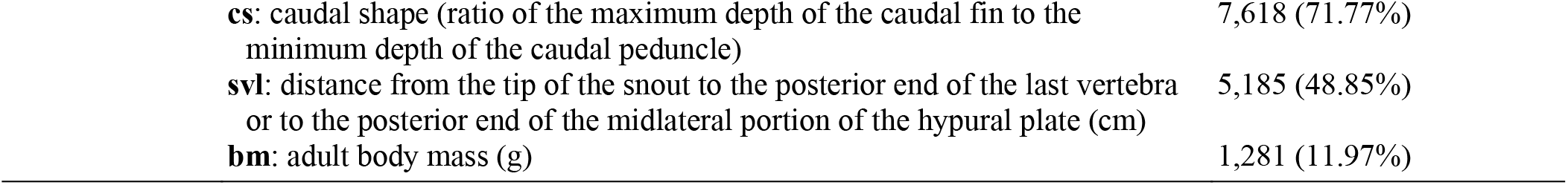
Functional traits considered in each group.

| Group | Trait | Number of species (% completeness) |
| --- | --- | --- |
| Plants | ph: Plant height (m) | 13,491 (34.77%) |
|  | ssd: Specific stem density (g/cm<sup>3</sup>) | 11,157 (28.76%) |
|  | la: Leaf area (mm<sup>2</sup>) | 12,791 (32.96%) |
|  | sla: Specific leaf area (mm<sup>2</sup>/mg) | 9,622 (24.80%) |
|  | ln: N content per unit leaf mass (mg/g) | 9,119 (23.50%) |
|  | sm: Seed mass (mg) | 23,096 (59.53%) |
| Mammals | ls: litter size (number of offspring per litter) | 3,511 (70.89%) |
|  | ly: number of litters per year | 2,146 (43.44%) |
|  | bm: adult body mass (g) | 4,651 (93.90%) |
|  | long: longevity (years) | 2,614 (52.78%) |
|  | gest: gestation length (days) | 2,220 (44.82%) |
|  | wea: weaning length (d) | 2,043 (41.25%) |
|  | fmat: time to reach female maturity (days) | 2,000 (40.38%) |
|  | svl: distance from the tip of the snout to the tail base (cm). | 3,921 (79.16%) |
| Birds | ls: clutch size (number of eggs) | 6,892 (70.31%) |
|  | ly: number of clutches per year | 1,784 (18.20%) |
|  | bm: adult body mass (g) | 9,532 (97.25%) |
|  | inc: incubation time (days) | 2,269 (23.15%) |
|  | long: longevity (years) | 1,672 (17.06%) |
|  | fa: fledging age (days) | 1,844 (18.81%) |
|  | em: egg mass (g) | 4,888 (49.87%) |
|  | svl: distance from the tip of the snout to the opening of the cloaca (cm) | 1,615 (16.48%) |
| Reptiles | ls: clutch size (number of eggs) | 2,675 (40.73%) |
|  | ly: number of clutches per year | 1,018 (15.50%) |
|  | bm: adult body mass (g) | 2,494 (37.98%) |
|  | inc: incubation time (days) | 1,369 (20.85%) |
|  | long: longevity (years) | 1,214 (18.49%) |
|  | svl: distance from the tip of the snout to the opening of the cloaca (cm) | 5,140 (78.27%) |
| Amphibians | am: age at maturity (years) | 384 (5.67%) |
|  | bs: body size (mm) | 5,227 (77.14%) |
|  | ls: maximum litter size (number of individuals) | 1,623 (23.95%) |
|  | os: offspring size (mm) | 1,330 (19.63%) |
| Freshwater fish | es: eye size (ratio of the diameter of the eye to the head depth) | 8,033 (75.68%) |
|  | ep: eye position (ratio of the centre of the eye to the bottom of the body to the body depth) | 8,033 (75.68%) |
|  | ms: mouth size (ratio of the length from snout to the corner of the mouth to the head depth) | 7,264 (68.44%) |
|  | mp: mouth position (ratio of the distance from the top of the mouth to the bottom of the body to the body depth) | 8,029 (75.65%) |
|  | elo: body elongation (ratio of the maximum body length to the maximum body depth) | 8,063 (75.97%) |
|  | wid: body lateral shape (ratio of the head depth at the vertical of eye to the maximum body depth) | 8,057 (75.91%) |
|  | ps: pectoral fin size (ratio of the length of the longest ray of the pectoral fin to body length) | 7,448 (70.17%) |
|  | pp: pectoral fin position (ratio of the distance between the upper insertion of the pectoral fin to the bottom of the body to body depth) | 7,999 (75.36%) |

|  |  |
| --- | --- |
| **cs:** caudal shape (ratio of the maximum depth of the caudal fin to the minimum depth of the caudal peduncle) | 7,618 (71.77%) |
| **svl:** distance from the tip of the snout to the posterior end of the last vertebra or to the posterior end of the midlateral portion of the hypural plate (cm) | 5,185 (48.85%) |
| **bm:** adult body mass (g) | 1,281 (11.97%) |

**Table S2.**
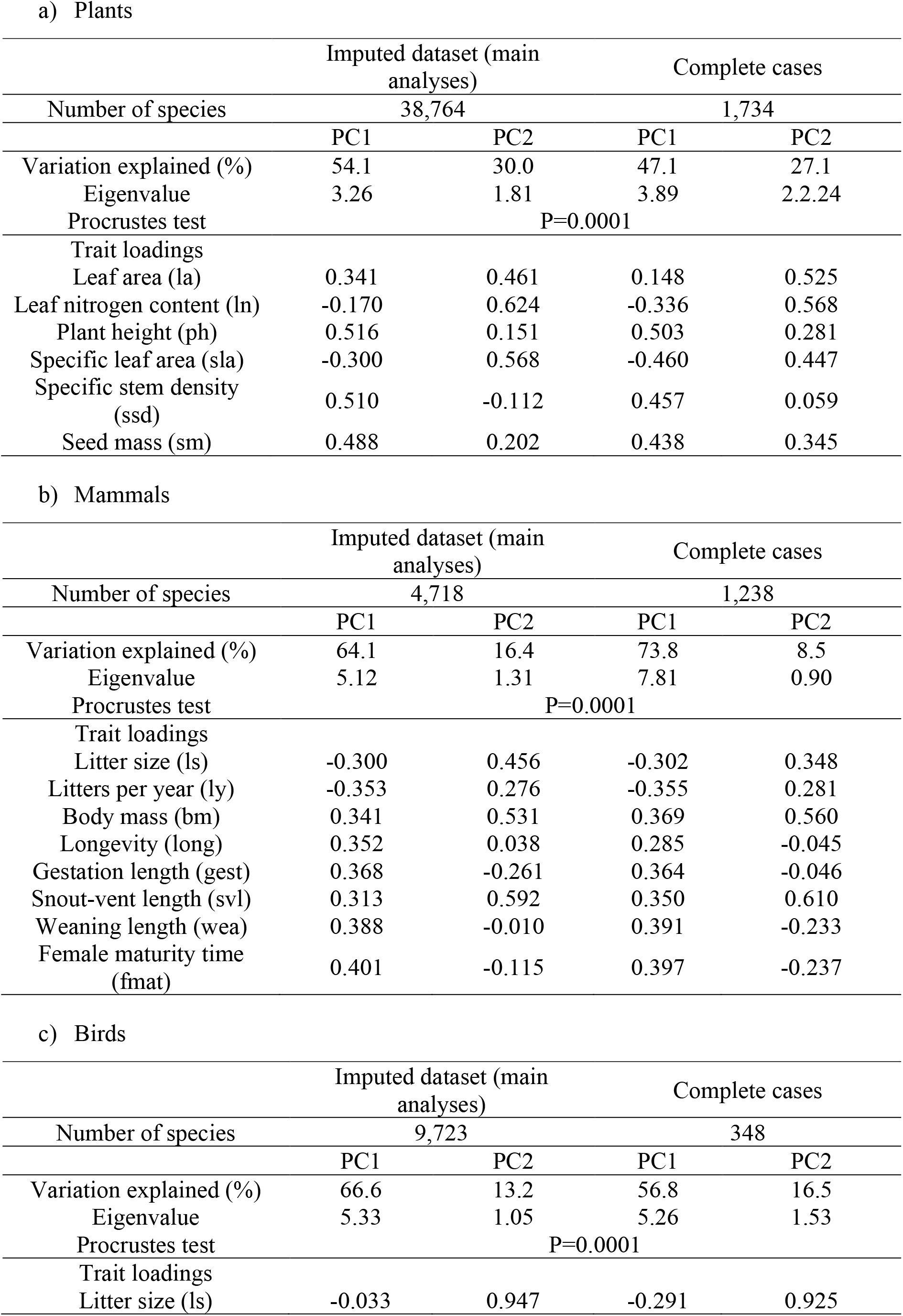
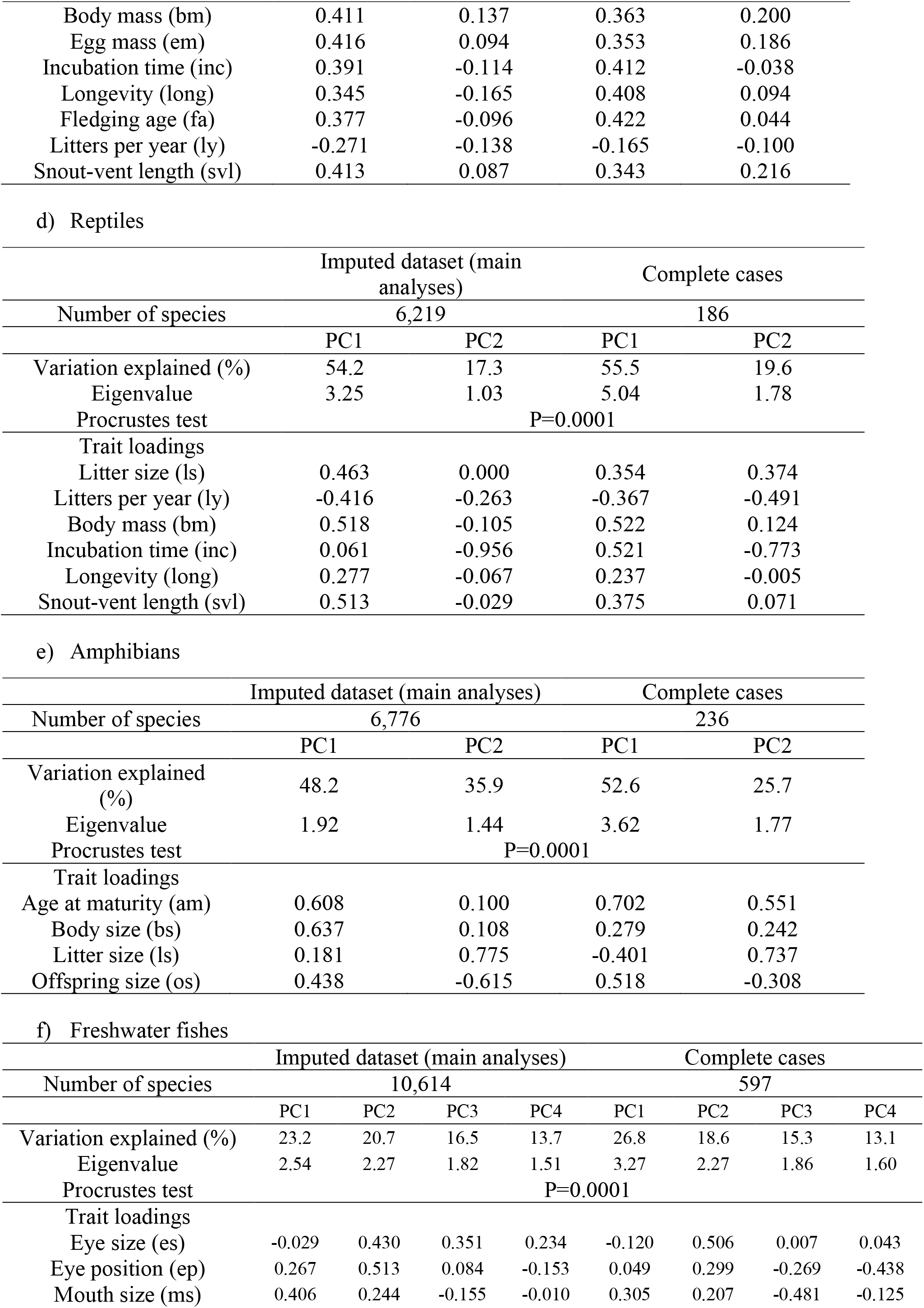
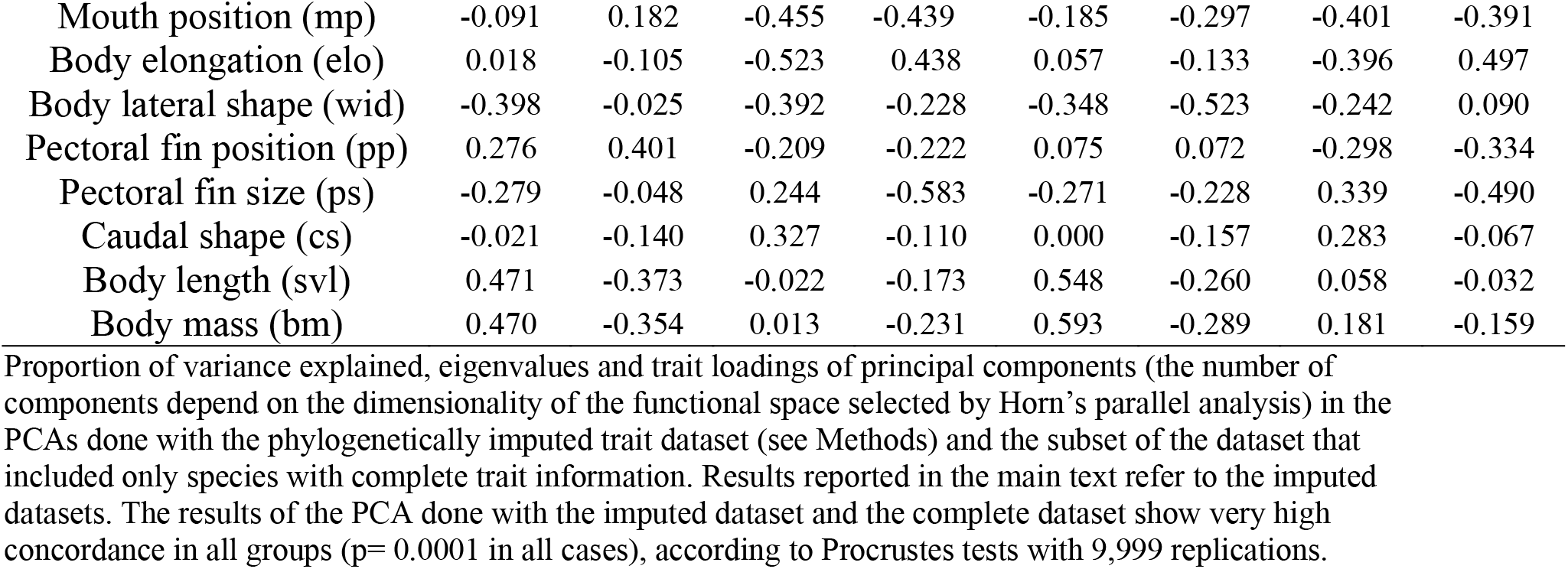
Principal component analysis (PCA) of trait data for the different taxonomic groups.

**Fig. S1.**
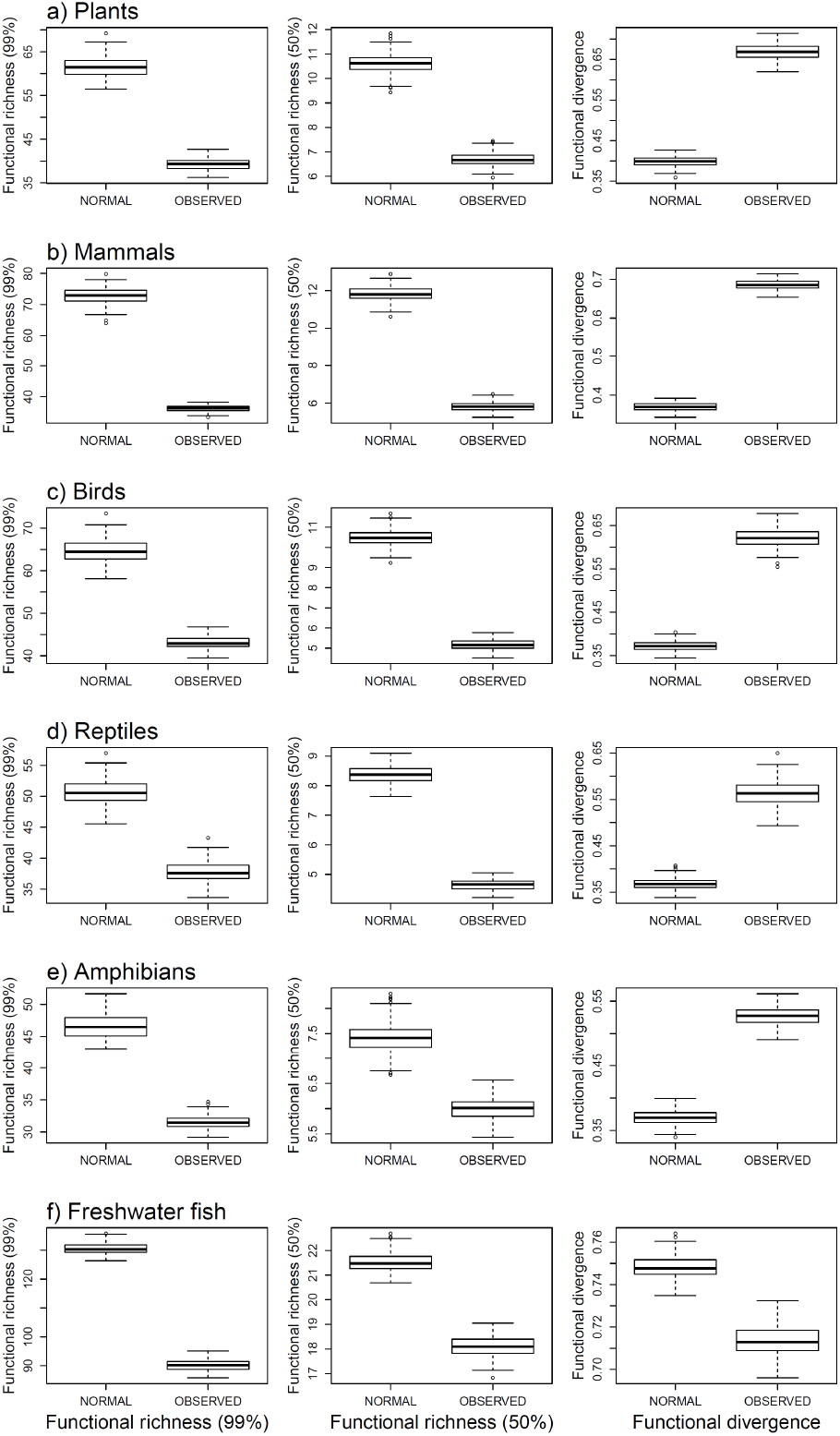
Comparison of functional structure metrics between observed spectra and equivalent multivariate normal distributions. For each taxonomic group considered (**a-f**), we drew 199 samples of 1,000 simulated species from multivariate normal distributions with the same mean and covariance matrix as the observed spectra. For each of these samples we estimated a TPD function and measured functional richness (amount of space occupied by the assemblages) at the and 99% and 50% quantile thresholds, and functional divergence (which indicates the degree to which the abundance of species in the functional trait space is distributed toward the extremes of its functional volume). Then, we drew 199 samples of 1,000 species from the observed global species pool of each taxonomic group and performed similar analyses. All three metrics showed distinctive patterns between the simulated multivariate normal (n=199 points) and the observed distributions (n=199). The functional volume occupied by the realised assemblages at the 99% and 50% thresholds were smaller than the corresponding multivariate normal distributions (two sided t-test, p<0.0001 in all cases), reflecting a high lumpiness in the distribution of species within the global functional spectra of all groups. Functional divergence of observed datasets was higher than that of the corresponding multivariate normal distributions in all groups except fishes (two-sided t-test, p<0.0001 in all cases), meaning that the most abundant species are close to the extremes of the distribution, except for fishes.

**Table S3.**
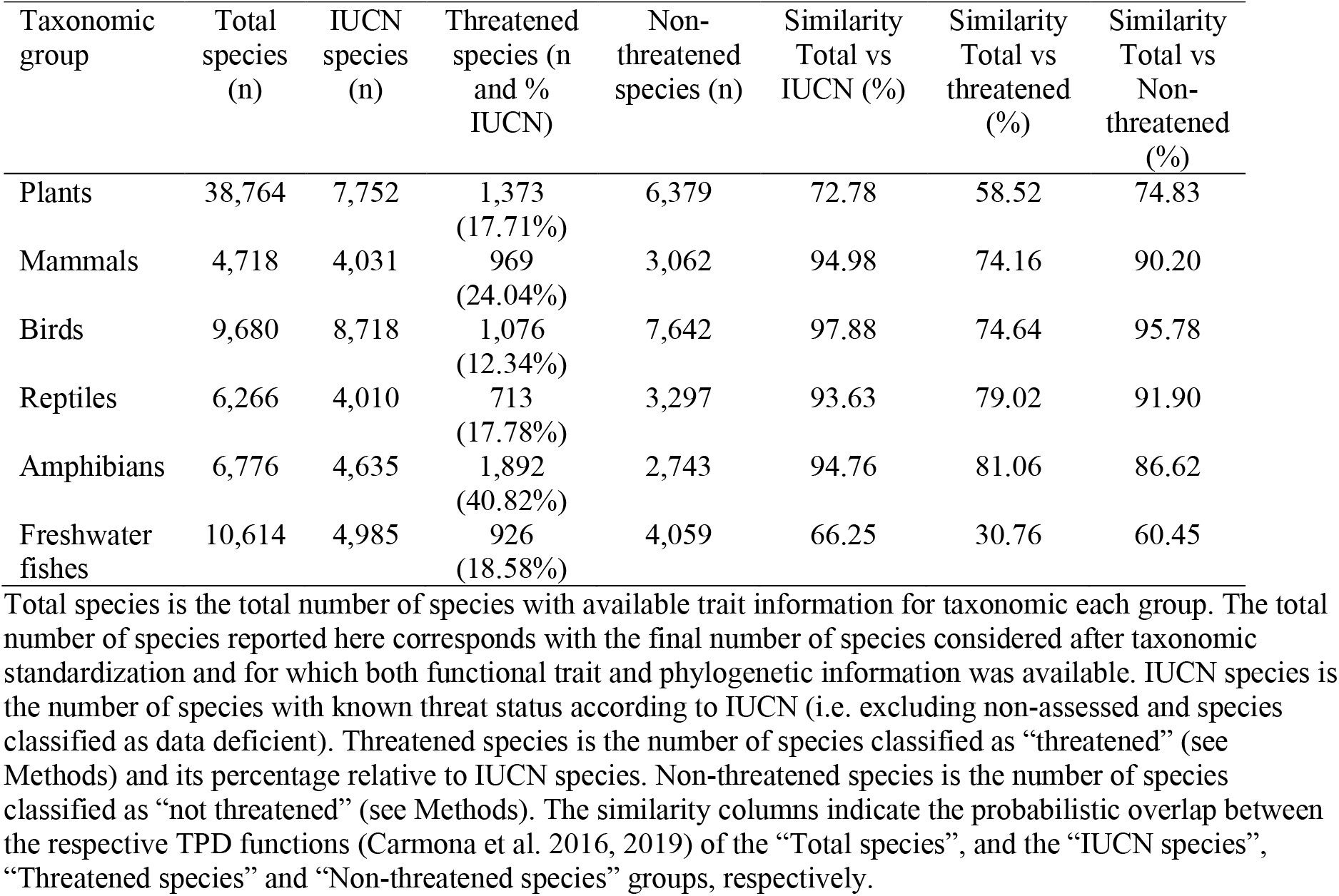
Similarity between functional spectra considering all species and functional spectra considering only species with known threat status according to IUCN.

| Taxonomic group | Total species (n) | IUCN species (n) | Threatened species (n and % IUCN) | Non-threatened species (n) | Similarity Total vs IUCN (%) | Similarity Total vs threatened (%) | Similarity Total vs Non-threatened (%) |
| --- | --- | --- | --- | --- | --- | --- | --- |
| Plants | 38,764 | 7,752 | 1,373 (17.71%) | 6,379 | 72.78 | 58.52 | 74.83 |
| Mammals | 4,718 | 4,031 | 969 (24.04%) | 3,062 | 94.98 | 74.16 | 90.20 |
| Birds | 9,680 | 8,718 | 1,076 (12.34%) | 7,642 | 97.88 | 74.64 | 95.78 |
| Reptiles | 6,266 | 4,010 | 713 (17.78%) | 3,297 | 93.63 | 79.02 | 91.90 |
| Amphibians | 6,776 | 4,635 | 1,892 (40.82%) | 2,743 | 94.76 | 81.06 | 86.62 |
| Freshwater fishes | 10,614 | 4,985 | 926 (18.58%) | 4,059 | 66.25 | 30.76 | 60.45 |

Total species is the total number of species with available trait information for taxonomic each group. The total number of species reported here corresponds with the final number of species considered after taxonomic standardization and for which both functional trait and phylogenetic information was available. IUCN species is the number of species with known threat status according to IUCN (i.e. excluding non-assessed and species classified as data deficient). Threatened species is the number of species classified as “threatened” (see Methods) and its percentage relative to IUCN species. Non-threatened species is the number of species classified as “not threatened” (see Methods). The similarity columns indicate the probabilistic overlap between the respective TPD functions (Carmona et al. 2016, 2019) of the “Total species”, and the “IUCN species”, “Threatened species” and “Non-threatened species” groups, respectively.

**Fig. S2.**
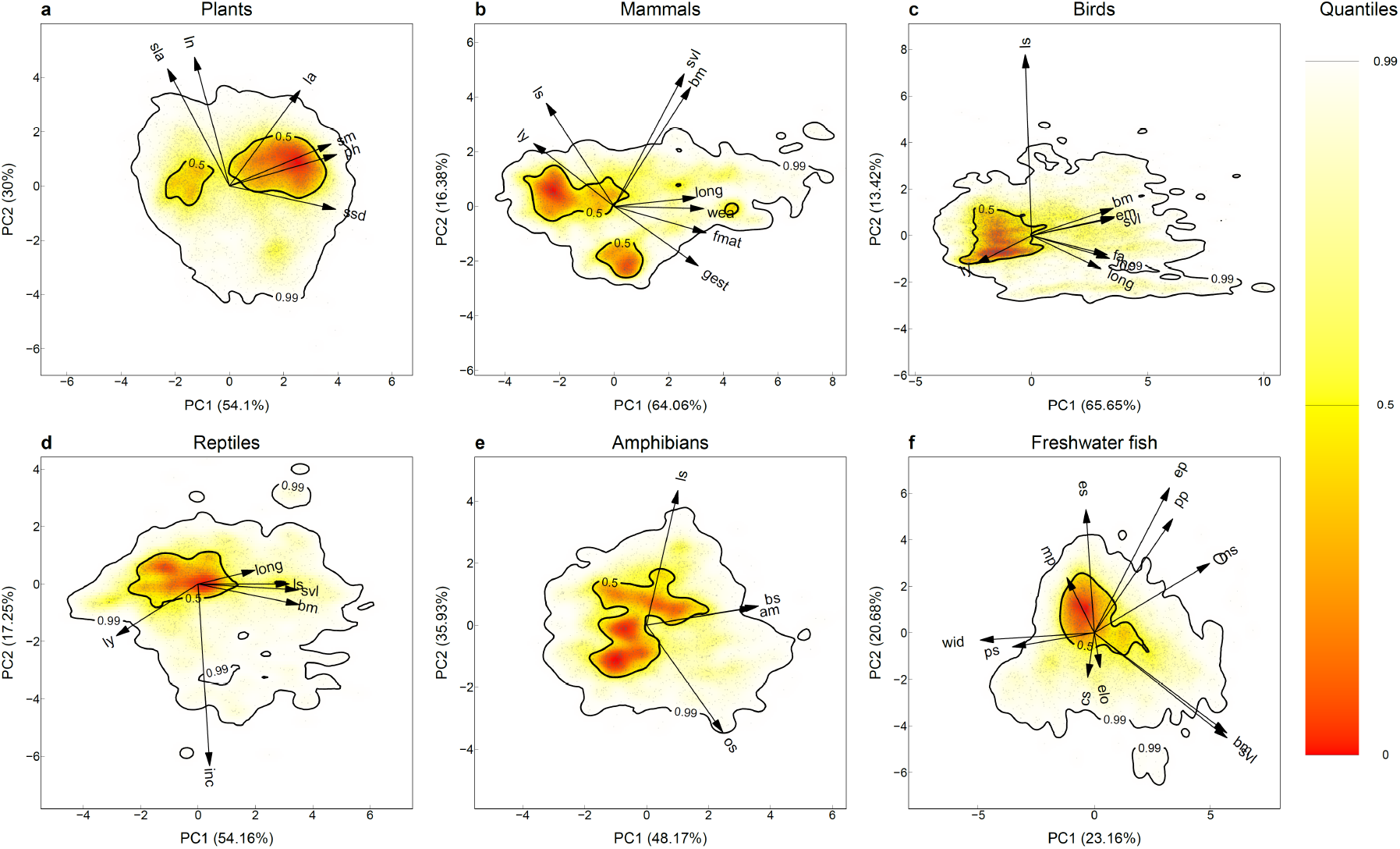
Global spectra of function of plants (A), mammals (B), birds (C), reptiles (D), amphibians (E) and freshwater fish (F) considering only species whose threat status has been assessed by IUCN. Probabilistic distributions of species assessed by IUCN on the spaces defined by the two first principal components of PCA considering different functional traits for each group. Arrows indicate the direction and weighting of each trait in the PCA (see Methods for description of each individual trait). The color gradient (red-yellow-white) depicts different density of species in the defined space (red areas are more densely populated). Arrows show the loadings of the considered traits in the resulting PCA. Thick contour lines indicate the 0.5 (hotspots, see main text) and 0.99 quantiles, and thinner ones indicate quantiles 0.1, 0.2, 0.3 and 0.4. Note the high correspondence (quantified in Extended Data Table 2) between the spectra shown here and those depicted in Fig. 1 in the main text, with the notable exception of plants.

**Fig. S3.**
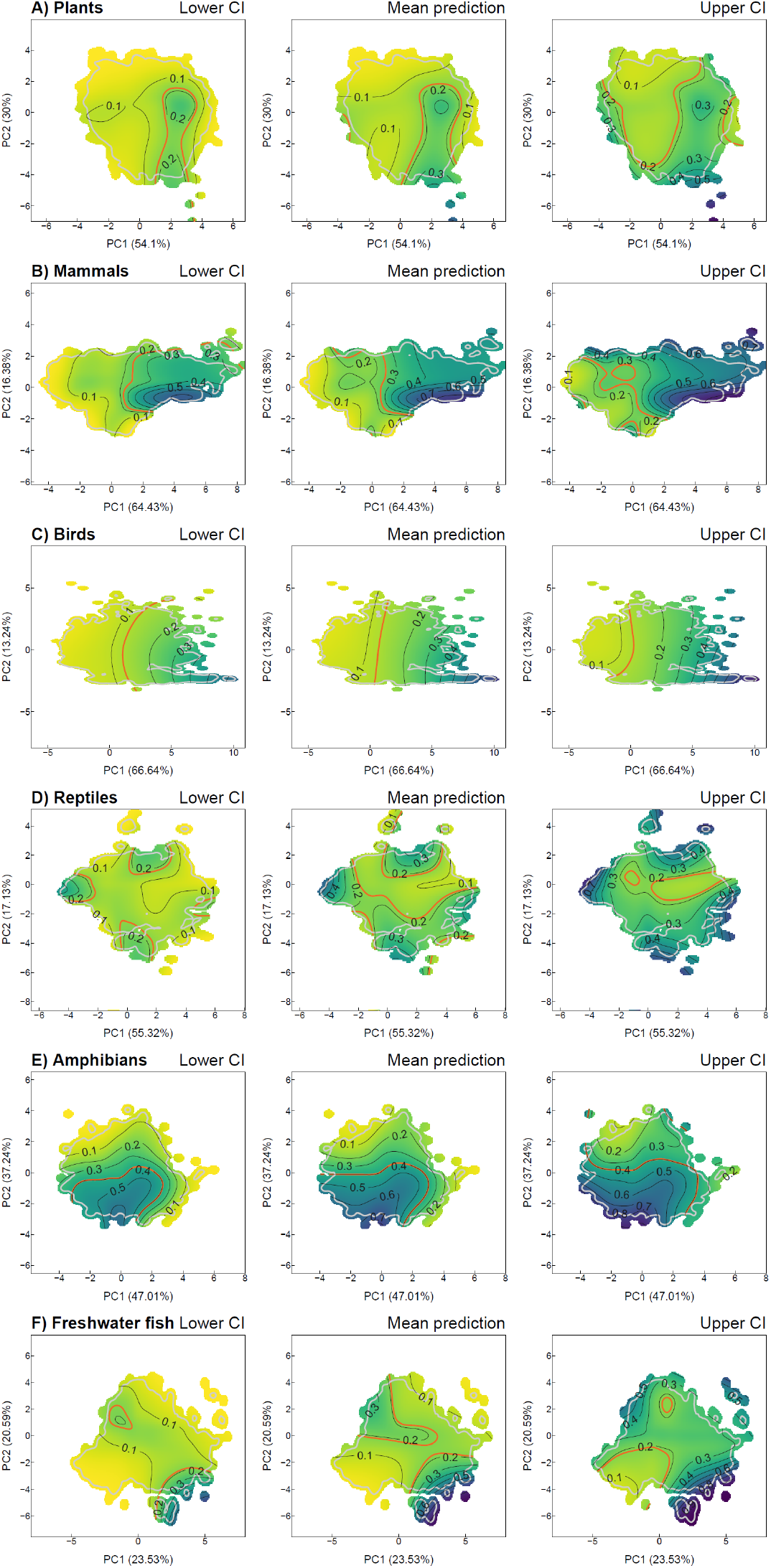
Uncertainty in the predictions of extinction threat in the functional spaces. The figure shows, for each group, the lower confidence interval of the mean (mean − 2 SE; left column), mean (center column), and upper confidence interval of the mean (mean + 2 SE; left column) of the probability of species being classified as threatened according to generalized additive models (GAM with binomial distribution) using the position of species in the functional space as predictors. Yellow tones indicate lower risk of extinction, whereas purple tones indicate high risk of extinction. For each group, the red contour line indicates the average threat probability (proportion of species classified as threatened). The grey line indicates the 0.99 quantile of the spectra of each group considering only species whose threat status is known.

**Movie S1.**

Animation showing the shifts in the plants functional spectrum after extinctions.

**Movie S2.**

Animation showing the shifts in the mammal functional spectrum after extinctions.

**Movie S3.**

Animation showing the shifts in the birds functional spectrum after extinctions.

**Movie S4.**

Animation showing the shifts in the reptiles functional spectrum after extinctions.

**Movie S5.**

Animation showing the shifts in the amphibians functional spectrum after extinctions.

**Movie S6.**

Animation showing the shifts in the freshwater fishes functional spectrum after extinctions.

